# Genomic offsets predict observed kelp declines and suggest benefits of assisted migration in the Northeast Pacific

**DOI:** 10.64898/2026.04.01.715974

**Authors:** Fernando Hernández, Jordan B. Bemmels, Samuel Starko, Loren H. Rieseberg, Gregory L. Owens

## Abstract

Kelp forests are widely distributed along temperate and polar coastlines worldwide and are among the world’s most productive and diverse marine ecosystems. Yet, due in part to ocean warming, they are declining in many parts of the world. While genomic tools can identify local adaptation and have shown potential to predict species’ responses to global change, these predictions have rarely been validated, hampering their widespread use in conservation practice. Here, we applied a seascape genomics approach to investigate environmental adaptation in the two main canopy-forming species of the Northeast Pacific, *Macrocystis tenuifolia* and *Nereocystis luetkeana*. We leveraged whole-genome sequences of 598 individuals across 94 sites along the British Columbia and Washington coasts, together with 37 environmental variables. Both species showed genomic signatures of local adaptation, with distinct environmental drivers shaping adaptation in each species despite their co-occurrence across much of the studied area. Using gradient forests, we modelled the genetic turnover across environmental gradients and estimated genomic offsets under projected environmental conditions. Genomic offsets differed greatly among regions in both species and help explain contrasting patterns of kelp vulnerability and resilience observed in local-scale studies. For *Macrocystis*, genomic offsets strongly predicted population declines observed to date, validating the link between genomic models and outcomes in the field and allowing us to translate genomic predictions into an ecologically meaningful metric: the risk of extirpation under global change. For *Nereocystis*, genomic offsets did not predict population declines well, suggesting that the environmental factors we modelled are not the primary drivers of *Nereocystis* persistence. Our models predict that assisted migration, even short-distance migration, could significantly attenuate kelp’s vulnerability to global change. Our results highlight the potential of seascape genomics to predict vulnerability of populations to global change and can inform conservation strategies to improve the resilience and sustainable management of these vulnerable ecosystems.

## INTRODUCTION

Kelp forests, formed by large brown macroalgae of the order Laminariales, are widely distributed along more than 1/3 of the world’s coastlines (Jayathilake & Costello 2021). They represent some of the most productive and diverse marine ecosystems, providing habitat for a wide range of species, including invertebrates, fishes, and other seaweeds of ecological and economic importance (Steneck et al., 2002, Teagle and Smale, 2018; Pessarrodona et al., 2022). However, these ecosystems are quickly disappearing in many parts of the world, mainly due to ocean warming and marine heatwaves (Vergés et al., 2016; Smale, 2020; Tait et al., 2021; Starko et al., 2022; Wernberg et al., 2024), though overgrazing by herbivores is also a major threat (Filbee-Dexter and Scheibling, 2014; Ling et al., 2015). The IPCC ranks kelp forests as the second most vulnerable coastal marine ecosystem to global change, only after coral reefs (Pörtner et al. 2019). Despite this general trend of decline, kelp ecosystems in some places remain stable or have even increased in abundance over the past several decades (Pfister et al., 2018; Smale, 2020; Mora-Soto et al., 2024; Starko et al., 2024). A clearer understanding of intraspecific and interspecific responses to environmental gradients is therefore critical to better predict responses to ongoing climate change and to inform conservation, management, and restoration strategies of kelp forest ecosystems.

In the Northeast Pacific, there are two main canopy-forming kelp species: the perennial giant kelp *Macrocystis tenuifolia* (hereafter *Macrocystis*) – until recently (Lindstrom 2023) included in *Macrocystis pyrifera* sensu lato (Demes et al. 2009) – and the annual bull kelp *Nereocystis luetkeana* (hereafter *Nereocystis*). As a result of past glaciation, the Northeast Pacific coastline is complex, featuring numerous fjords, islands, and shorelines of varying wave exposure levels, which creates a mosaic of microclimates with high variability in key ocean variables (e.g., temperature, salinity, nutrients) over relatively short distances (Mora-Soto et al., 2024; Starko et al., 2024; Man et al., 2025). A recent genetic study revealed strong genetic structure associated with geography in both species (Bemmels et al., 2025). However, the extent to which populations are locally adapted to these environmental gradients and the amount of intraspecific genetic diversity for adaptive loci remain unknown. Importantly, complex environmental heterogeneity in the Northeast Pacific decouples environmental gradients from simple geographic proxies such as latitude (Mora-Soto et al., 2024; Starko et al., 2024; Gendall et al. 2025). In many systems, spatial and environmental heterogeneity are often strongly confounded, making it challenging to tease apart isolation-by-distance from true signatures of local adaptation (e.g., Vranken et al. 2021; Wood et al. 2021; Minne et al., 2025). In contrast, localities at the same latitude can vary substantially in environmental conditions in the Northeast Pacific (Mora-Soto et al., 2024; Starko et al., 2024; Gendall et al. 2025), potentially enabling more robust inferences about genotype-environment relationships.

As both species have recently experienced strong declines in some parts of the Northeast Pacific (e.g., Mora-Soto et al., 2024; Starko et al., 2024; Gendall et al. 2025), interest in kelp forest restoration is rapidly growing (Wood et al., 2024a; Dykman et al., 2025). To preserve genetic integrity of restored sites, restoration best-practices that involve moving individuals from one location to another often recommend using local source populations whenever possible (Bucharova et al., 2017). When suitable local sources are not available, population genetic structure has also been used to choose donor sources. This has been done successfully in other kelp species (Wood et al., 2020) and experimentally in bull kelp in British Columbia (BC) (L. Dykman and J. Baum, personal communication). In the absence of detailed knowledge about local adaptation or population genetic structure, government agencies in Alaska (Gruenthal and Habicht, 2022), BC (McConnell et al. 2024), and Washington (Cui, 2023) have adopted informal but conservative recommendations that kelp be transferred no further than 50 km from its geographic provenance for restoration and aquaculture. However, restoration with local germplasm is unlikely to be successful if environmental conditions have already shifted away from the historical conditions to which local genotypes were adapted (Houde et al., 2015; Coleman et al., 2020).

To address such concerns, methods known as genomic offsets (GOs) have recently been developed to predict population responses to environmental variation in space and time (Fitzpatrick and Keller, 2015; Capblancq et al., 2020). Genomic offset statistics rely on genotype-environment association (GEA) analyses to model changes in allele frequencies along current environmental gradients. Then these models are used to predict allele frequencies from environmental features and evaluate differences in predicted allele frequencies at pairs of temporal or spatial points (Capblancq et al., 2020; Gougherty et al., 2021). Higher genomic offsets suggest that a population’s genomic composition will be less suited to the given environment in a different location or different time period, though suitability will depend on the relationship between genomic offsets and fitness (Lachmuth et al., 2024; Lotterhos, 2024). This approach offers a powerful toolkit to inform decisions about selection of the optimal genotypes for restoration and the identification of areas with higher or lower vulnerability under changing climates.

Though little is known about local adaptation in most marine species in general (Layton et al. 2024), previous studies in other kelp and seaweeds have calculated genomic offsets and identified environmental gradients and candidate loci involved in adaptation (Vranken et al. 2021; Wood et al. 2021; Minne et al. 2025), highlighting how this approach has the potential to inform restoration strategies in kelp (Coleman et al., 2020). GEA analysis has also been recently applied to *Nereocystis* and used to identify candidate source populations for genetic rescue (Abbott et al. 2026); however, GEA was applied on only a very small portion of this species’ range (Puget Sound, Washington) and genomic offsets were not explicitly calculated. Furthermore, as GEA analyses rely on statistical associations between genetic variants and environmental variables, their predictions are rarely experimentally tested (but see Fitzpatrick et al., 2021; Luo et al., 2025; Verrico et al., 2026; reviewed in Fitzpatrick et al., 2026). In species such as kelp for which long-term ecological monitoring datasets are available (e.g., Starko et al., 2024), comparing genomic offsets to ecological performance could provide a powerful approach to validating GEA models and translating genomic offsets into quantified predictions about ecological outcomes.

Here, we investigate the genetic bases of environmental adaptation in the two main canopy-forming species from the coasts of British Columbia and Washington: *Macrocystis* and *Nereocystis*. Our aims are to 1) identify the most relevant environmental gradients for each species; 2) predict the vulnerability of kelp forests to global change under various migration scenarios; and 3) validate our predictions with contemporary kelp persistence records. Together, these approaches provide a framework for linking environmental adaptation to future vulnerability and informing conservation strategies in Northeast Pacific kelp forests.

## METHODS

### Genetic variation and population structure

We obtained initial pools of 10,800,722 and 16,527,403 high-quality SNPs from re-sequencing 191 and 404 individuals of *Macrocystis* and *Nereocystis*, respectively. These individuals were collected from 33 and 61 sites (each with at least three individuals; hereafter “populations”). These initial SNP pools were previously generated by Bemmels et al. (2025) and were filtered to exclude first-degree relatives. For each species, we further processed the initial pools to generate two filtered SNP datasets: 1) the full dataset filtered to a minimum minor allele frequency (MAF) of 0.05, excluding only redundant fully linked loci (r^2^ < 0.99), which gave 375,110 and 1,228,456 SNPs for *Macrocystis* and *Nereocystis*, respectively; and 2) a low linkage disequilibrium (LD) dataset also filtered to a minimum MAF of 0.05 but including only SNPs with low LD (r^2^ < 0.2 using sliding windows of 50 kb and 10 kb steps), which gave 43,933 and 116,707 SNPs for *Macrocystis* and *Nereocystis*, respectively. Using the SNPs generated in these datasets, we calculated the allele frequency for each SNP and population and used the population-level allele frequency data for all subsequent analyses. The full datasets were used for genotype-environment associations, while the low LD datasets were used for principal component analysis.

### Environmental characterization of kelp forest habitats

To characterize the environmental variation experienced by kelp, we retrieved 37 environmental variables from the Northeastern Pacific Canadian Ocean Ecosystem Model (NEP36-CanOE) Climate Projections database (Holdsworth et al. 2021). These variables included mean, minimum, and maximum values for sea surface temperature, salinity, nutrient availability, and primary productivity (Table S1). For each variable, we obtained modelled historical values (1986-2005) as well as simulations for the period 2046-2065 under two global warming scenarios: moderate mitigation (RCP4.5) and no mitigation (RCP8.5). The NEP36-CanOE environmental variables represent outputs of an ocean circulation model for the historical period (plus climate change projections for future time periods), rather than observed data points associated with actual kelp populations and thus represent a source of uncertainty incorporated into our genomic models. We also included fetch (a proxy for wave exposure) as wave exposure is known to impact kelp habitat suitability (e.g., Mora-Soto et al. 2024; Graham et al. 1997). Fetch was defined as the total distance from a focal ocean point to land (up to a maximum of 200 km) summed across 72 compass bearings separated by 5° each (Gregr et al. 2018) and was calculated using custom *R* scripts as the mean fetch across 100 evenly spaced points per pixel of a 1/36°-resolution grid (excluding points that fell on land). The coastline map used in fetch calculations was a composite of source maps for BC and Washington (GeoBranch BC 2002) and Alaska (Alaska DNR 2021), with both source maps at 1:250,000 resolution.

January and July sea surface values of environmental variables (except fetch, for which only a single value applies) were extracted for a grid dataset at 1/36° resolution covering the range of distribution of the two species, comprising 4,447 and 7,479 observations for *Macrocystis* and *Nereocystis*, respectively. January and July were selected as proxies for winter and summer environments, respectively. The spatial masking for each of the species was approximated based on species distribution data from the BC Marine Conservation Atlas (BC Marine Conservation Analysis 2011). Using only the historical values, we ran a pairwise correlation analysis to filter out environmental variables with a Spearman correlation coefficient (r) > |0.8|. In general, January and July values of each variable showed low correlation, only Nitrate concentration in January (NO_3__01) and July (NO_3__07) passed the threshold, and we retained NO_3__07. For each month (January or July), the minimum, mean, and maximum values of each variable showed strong correlation, so we retained the mean values. Among environmental variables, we retained sea water practical salinity (salt_01 and salt_07) over total alkalinity concentration, sigma potential density, and dissolved inorganic concentration, and pH over air-sea CO_2_ flux. The final set comprised 14 environmental variables (Table 1 and S1). Principal component analysis using the 14 environmental variables (zero centered) from observed and sampled sites was performed using the R function *prcomp*.

**Table 1.** Final set of environmental variables used in this study. Mean, minimum, and maximum values of sampled populations are shown. * Following Gregr et al. (2018).

| Variable | Description | Unit | Source | Mean | Minimum | Maximum |
|---|---|---|---|---|---|---|
| Fetch | log10 mean Sum Fetch | log10 (m) | This study* | 5.8 | 4.6 | 6.8 |
| NO<sub>3</sub>_07 | Nitrate concentration in July | mmol m-3 | NEP36-CanOE | 3.4 | 5.4E-07 | 13.7 |
| O<sub>2</sub>_01 | Dissolved oxygen concentration in January | mmol m-3 | NEP36-CanOE | 305 | 276 | 346 |
| O<sub>2</sub>_07 | Dissolved oxygen concentration in July | mmol m-3 | NEP36-CanOE | 284 | 256 | 309 |
| omega_a_01 | Aragonite saturation state in January |  | NEP36-CanOE | 1.7 | 1.1 | 1.9 |
| omega_a_07 | Aragonite saturation state in July |  | NEP36-CanOE | 2.1 | 1.2 | 2.5 |
| PH_01 | pH in January |  | NEP36-CanOE | 8.1 | 8.1 | 8.2 |
| PH_07 | pH in July |  | NEP36-CanOE | 8.1 | 8.0 | 8.2 |
| salt_01 | Sea water practical salinity in January | PSU | NEP36-CanOE | 28.3 | 18.3 | 31.1 |
| salt_07 | Sea water practical salinity in July | PSU | NEP36-CanOE | 28.2 | 14.6 | 31.8 |
| temp_01 | Mean sea surface temperature in January | Deg C | NEP36-CanOE | 7.2 | 3.3 | 9.4 |
| temp_07 | Mean sea surface temperature in July | Deg C | NEP36-CanOE | 13.6 | 10.2 | 17.6 |
| TPP_01 | Total primary production in January | mol m-3 s-1 | NEP36-CanOE | 1.0E-08 | 2.2E-09 | 2.2E-08 |
| TPP_07 | Total primary production in July | mol m-3 s-1 | NEP36-CanOE | 6.7E-08 | 8.0E-11 | 1.3E-07 |

### Identification of genomic regions associated with local environmental adaptation

We used latent factor mixed models (LFMM) with ridge penalty as implemented in the *lfmm* version 2.0 R package (Caye et al., 2019) to test for associations between genotypes (allele frequencies) and each of the 14 environmental variables while accounting for population structure. In LFMM, the genotype matrix is the response variable in a regression mixed model, the environmental variables are the explanatory variables introduced as fixed effects, and population structure is modeled using latent factors. Latent factors are unobserved variables estimated from the genotype matrix itself that capture covariance patterns shared genome wide. Then, the effect of environmental variables is tested on the residual component, largely reducing the rate of false positives (Caye et al., 2019). The number of latent factors is chosen based on estimates of population structure (in our case using principal component analysis) and represents the most likely number of ancestral genetic groups. We used three latent factors for both species, as this is the most likely number of ancestral genetic groups (Fig. S1). P-values were calibrated by calculating the genomic-inflation factor, an estimate of variance of z-scores, using the *lfmm_test* function (Caye et al., 2019).

Then, calibrated p-values were combined using a windows-based approach, Weighted-Z Analysis (WZA) (Booker et al., 2024). This approach combines information from multiple linked sites within *a-priori* defined windows to identify regions (instead of individual SNPs) associated with environmental variables. The approach uses expected heterozygosity (estimated from minimum allele frequencies) to construct weights for SNPs, as sites with higher expected heterozygosity carry more information about the demographic history of populations (Booker et al., 2024). The window size was set to 10 kb, and only windows with more than five SNPs were considered. A strict Bonferroni correction (p < 0.05/number of windows) was used for setting the significance threshold. Although windows chosen by WZA have higher average Z scores, not all sites within a window are necessarily adaptive. Therefore, for all SNPs within significant windows, we individually calculated pairwise correlations between allele frequency and each of the environmental variables (environment), latitude and longitude (geography), and the first four principal components from the PCA (population structure) and retained only those SNPs whose correlation was stronger with any environmental variable than geography and population structure variables. Hereafter, we refer to these SNPs as ‘adaptive’.

To validate our set of adaptive SNPs and ensure that patterns were likely due to environmental adaptation rather than statistical artifacts, we generated, for comparison, a set of neutral SNPs by randomly selecting a similar number of windows not associated with any environmental variable (all p-values > 0.1). Mantel tests and gradient forest analyses were performed for neutral and adaptive SNPs. Mantel tests were used to test for isolation by distance (IBD) and isolation by environment (IBE) under the expectation that adaptive SNPs should have greater IBE. For Mantel tests, environmental distances were calculated as Euclidean distances among the 14 environmental variables, while geographic distances were calculated as the minimum ocean distance (i.e., excluding land) between populations and were previously generated by Bemmels et al. (2025). Gradient forest analyses were used to inspect the rate of allele turnover along environmental gradients for neutral and adaptive SNPs. Adaptive SNPs are expected to show stronger and faster turnover rates than neutral SNPs due to the combined forces of selection and demography. These analyses were performed using the R packages *vegan* version 2.6.4 (Oksanen et al. 2022) and *gradientForest* version 0.1.37 (Ellis et al. 2012).

### Genomic offsets to predict vulnerability to climate change

By using the set of adaptive SNPs and the 14 environmental variables, we investigated spatial and temporal patterns of maladaptation to future climate change for both species. We used gradient forests (GF) (Ellis et al. 2012) to model the genetic turnover along present environmental gradients and predict genomic offsets to projected future environments. We estimated three complementary genomic offset (GO) variables, each quantifying the degree of mismatch between current genetic composition and future environmental conditions. First, we modelled local genomic offset, which quantifies *in situ* mismatch and does not allow migration of genotypes. Next, we modelled forward and reverse genomic offset (as defined by Gougherty et al., 2021), which allow migration of genotypes across space but using different approaches. To aid understanding, in this study, we refer to forward and reverse genomic offsets as Population GO and Location GO, respectively. Population GO predicts how the current population genotype will fare in the future, assuming it can move to the most optimal spot available across the predefined mapping space. A low Population GO indicates that the genotype has a suitable future home, while a high Population GO means that the genotype will be less adapted in the future than at present, regardless of where it moves. Location GO asks whether any current populations have a genetic composition that is well adapted to the future environmental conditions of that location. A low Location GO score means that there are populations currently available with a genomic composition that is well adapted to the location’s future climate. A high Location GO indicates that no current genotype will be adapted to the future climate at the location. Under this framework, local GO fixes genotypes in place, Population GO fixes the genetics of a population but allows its location to change, while Location GO fixes location, but allows genotypes to disperse to this location from any current population.

To explore how Population GO and Location GO would be impacted by different kelp management strategies, we considered two additional versions of the GOs. The first version limited migration to a set maximum distance of 50 km, in line with current guidelines for the maximum distance that kelp can be transplanted in Alaska, BC, and Washington (Gruenthal and Habicht 2022; Cui 2023; McConnell et al. 2024). In this version of the GO analysis, Population GO_50 km_ asks how well the genotype at a location would do in the future, assuming it could only migrate or be transferred up to 50 km; Location GO_50 km_ asks if there are suitable genotypes to colonize its future climate within 50 km. The second additional version of the GO analysis is similar but instead of 50 km it considers migration within pre-defined geographic regions for each species (Population GO_REGION_ and Location GO_REGION_) (Fig. 1A). These regions were largely delineated based on neutral population genetic structure (Bemmels et al. 2025), with minor adjustments to reflect geographic boundaries and areas of conservation importance.

**Figure 1.**
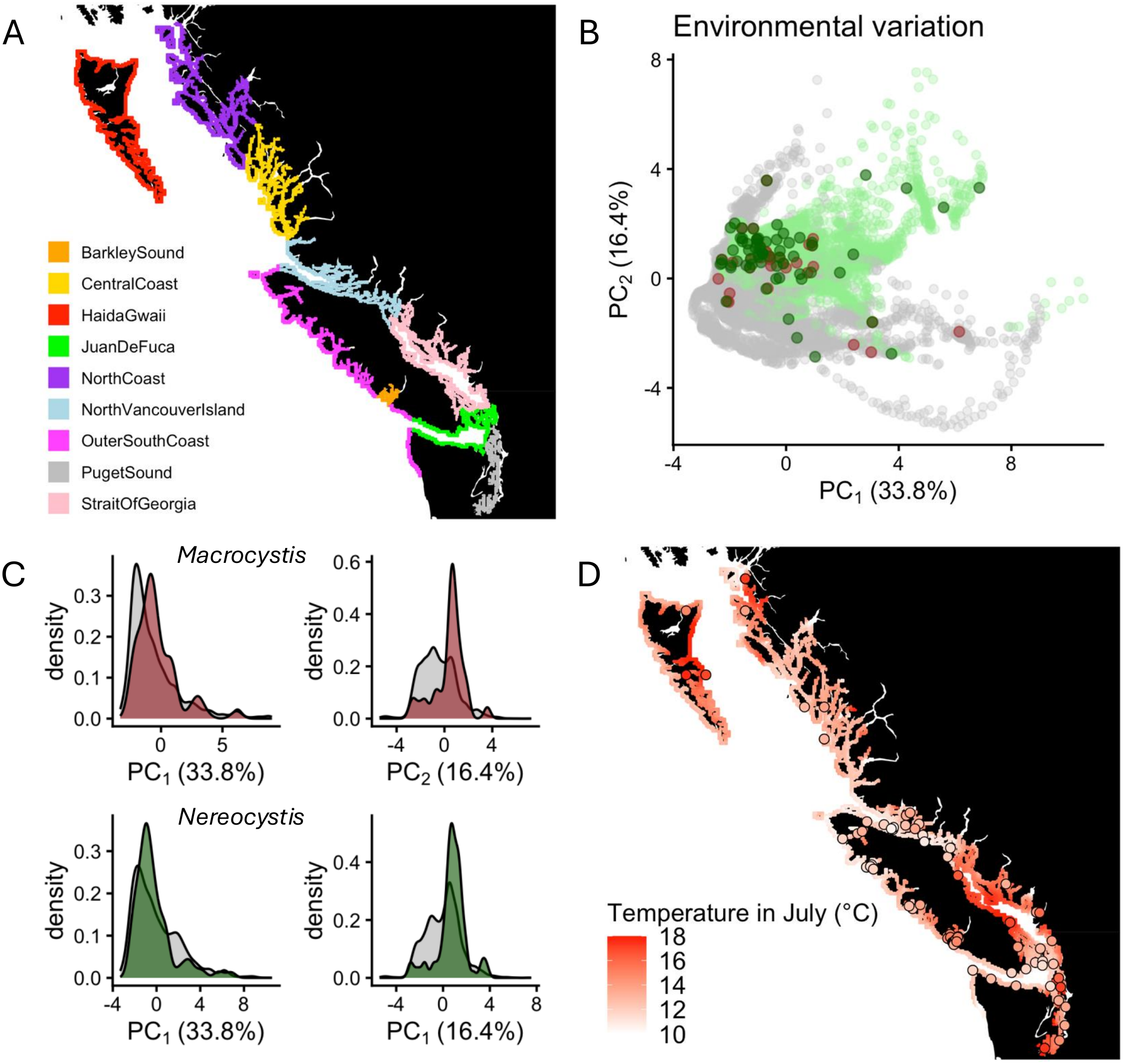
Environmental variation and sampled sites. A: Pre-defined geographic regions used in this study. *Nereocystis* is present in all regions, while *Macrocystis* is present in all regions but Puget Sound and Strait of Georgia. B: Principal component analysis using 14 environmental variables with low correlation (r < |0.8|). Background gray and green dots correspond to sites where both species and only *Nereocystis* are potentially present (Puget Sound and Strait of Georgia), respectively, based on their broad-scale biogeographic distributions. Brown and dark green main dots represent sampled populations used in this study for *Macrocystis* and *Nereocystis*, respectively. C: Density plots showing the environmental distribution of occurrence (gray) and sampled (brown or dark green) sites for *Macrocystis* (top panels) and *Nereocystis* (bottom panels). D: Mean sea surface temperature in July showing the complex geographic pattern of environmental variation along the BC coastline. Circles corresponded to sampled sites, which may be plotted rounded to the nearest 0.5° to protect information about culturally sensitive sites (see Biocultural Notice in Bemmels et al., 2025).

### Validation of genomic offsets

To validate our genomic offset estimates, we used kelp abundance data from Starko et al. (2024) for each species and asked whether sites with extirpated populations show higher genomic offsets than sites with stable populations. We registered presence-absence at two timepoints: T1 (1997-2007) and T2 (2018-2021). The timepoints were separated by 14 to 27 years (median = 14 years for both species). Sites where the species was present at T1 were classified as ‘stable’ if the species was present at both timepoints or ’loss’ if it was absent at T2. Sites where the species was absent at T1 were not included in the analyses. Then, for each species, we calculated local genomic offsets for 2020 as above. Values of environmental variables for 2020 were estimated as the average between historical (1986-2005) and projected values for the period 2046-2065 under the two projected scenarios (RCP.4.5 and RCP8.5). Finally, for each site with presence-absence data (stable or loss), we extracted the genomic offset of the closest point in the grid (mean distance = 1.07 and 1.04 km for *Macrocystis* and *Nereocystis*, respectively), which gave us 295 and 447 sites with stable or loss data from 83 and 97 points with genomic offset estimates for *Macrocystis* and *Nereocystis*, respectively. Weighted logistic regressions were used to model the probability (hereafter risk) of extirpation as a function of the estimated genomic offset for 2020 and compared to a null model with temp_07 as predictor, using AIC and McFadden-pseudo-r^2^ as metrics. Although we ran the model with each of the environmental variables as predictor, we chose temp_07 as the null model because in both species summer temperatures have been associated to population declines in the Northeast Pacific coastline (e.g., Mora-Soto et al., 2024; Starko et al., 2024).

For *Macrocystis*, we estimated donor and recipient importance (DI and RI, respectively), which quantifies the percentage of potential suitable locations in the future for each current genotype and the percentage of potential suitable current genotypes for each future location, respectively (Lachmuth et al., 2024). A location with high DI is expected to be suitable in many different locations under future climates, while a location with high RI has many possible donors that are expected to be suitable. These statistics, by counting the number of suitable locations rather than a single best-match, are more robust for estimating the overall value of populations and locations as potential sources and recipients, respectively (Lachmuth et al., 2024). To avoid an arbitrary genomic offset threshold, suitable populations/locations were those with scaled offset values below the z-score threshold, which corresponds to a predicted risk of extirpation of 50%. For these analyses, we analyzed only *Macrocystis* because model fit was substantially better for *Macrocystis* than for *Nereocystis*.

## RESULTS

### Environmental characterization of kelp forest habitats

In the environmental PCA, the first two axes explained 51% of variation (Fig. 1B). The environmental distribution of both species largely overlaps (Fig. 1C), and our sampled sites span most of the environmental variation found in the region (Fig. 1B, C). Although our sampling spanned over seven degrees latitude, temperature did not follow latitude (Fig. 1D). Some of the northern regions, for example, inner Haida Gwaii, had higher summer temperatures than many southern areas. Also, most locations in the Strait of Georgia had higher summer temperatures than locations from similar latitudes in the Outer South Coast.

### Identification of genomic variants associated with local environmental adaptation

We identified 41 and 77 10 kb windows significantly associated with environmental variation in *Macrocystis* and *Nereocystis*, respectively. Associated windows were widely distributed across the genome, containing 1,914 (5-120 per window) and 2,910 (9-102 per window) SNPs in *Macrocystis* and *Nereocystis*, respectively (Fig. 2A). Most environmental variables, all but sea water practical salinity in January and July (salt_01 and salt_07, respectively), were significantly associated with at least one genomic region (Fig. 2B). For *Macrocystis*, aragonite saturation state in January (omega_01), followed by dissolved oxygen concentration in January (O_2__01) and nitrate concentration in July (NO_3__07), showed the most associations while for *Nereocystis*, fetch (a proxy for wave exposure), followed by mean sea surface temperature in January (temp_01) and July (temp_07), showed the most associations. Further selecting only SNPs that had a stronger association with environment than population structure or geography further restricted the datasets to 944 and 1,590 SNPs, hereafter ‘adaptive’ SNPs. For comparison, we used 1,000 and 1,600 SNPs from 41 and 77 randomly chosen windows not associated with any environmental variable (all p-values > 0.1) for *Macrocystis* and *Nereocystis*, respectively. Hereafter, we refer to this set of SNPs as ‘neutral’.

**Figure 2.**
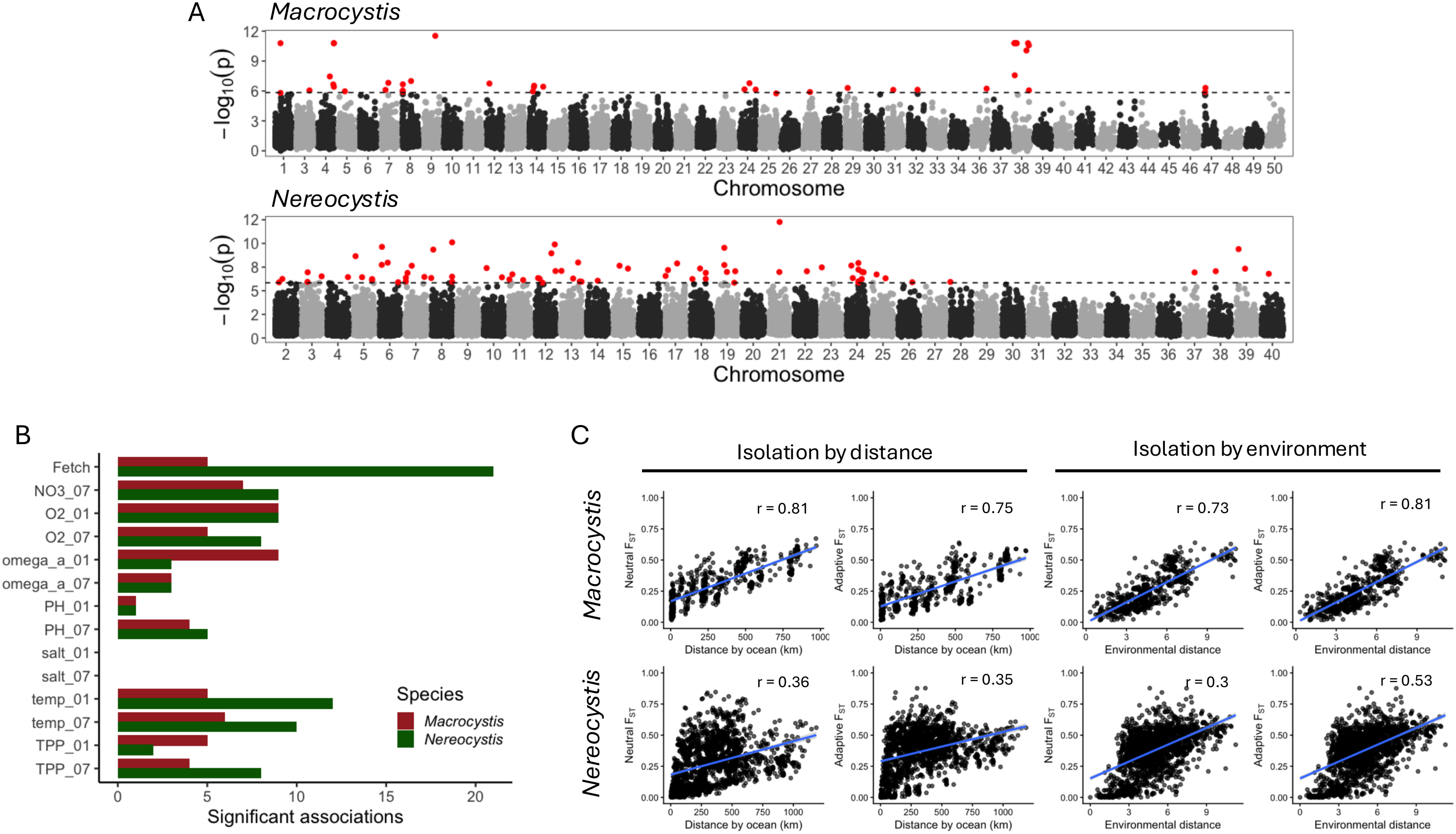
Genetic architecture of environmental adaptation. A: Manhattan plots showing the association between 10 kb windows and environmental variables; only the top association across variables is shown for each window. B: Number of significant associations found for each environmental variable. C: Isolation by distance (IBD) and isolation by environment (IBE) for neutral and adaptive loci for *Macrocystis* (top panels) and *Nereocystis* (bottom panels).

We used two complementary approaches to compare adaptive and neutral sets of SNPs. First, we performed Mantel tests to evaluate patterns of isolation by distance (IBD) and isolation by environment (IBE). IBD was substantially stronger for *Macrocystis* than for *Nereocystis* (r^2^ = 0.81 and 0.36, respectively; Fig 2C), and it was slightly stronger for neutral than adaptive SNPs (r^2^ = 0.81 vs. 0.75 and 0.36 vs. 0.35 for *Macrocystis* and *Nereocystis*, respectively; Fig. 2C). In contrast, IBE was stronger for adaptive than neutral loci in both species (r^2^ = 0.81 vs. 0.73 and 0.53 vs. 0.3 for *Macrocystis* and *Nereocystis*, respectively; Fig. 2C), providing validating evidence for the selection of adaptive loci. When IBD and IBE were analyzed together using multiple regression, both IBD and IBE were significant in both species and sets of SNPs (r^2^ = 0.70 and 0.72 and 0.16 and 0.30 for neutral and adaptive SNPs in *Macrocystis* and *Nereocystis*, respectively). Finally, we used a machine learning approach (gradient forest, GF), to show that the allele frequency turnover was stronger and more rapid for adaptive than neutral SNPs in both species (Fig. 3). Together, these complementary analyses indicate that our set of adaptive loci are more strongly structured by environmental variation than by geographic distance alone and therefore likely reflect meaningful signals of local adaptation.

**Figure 3.**
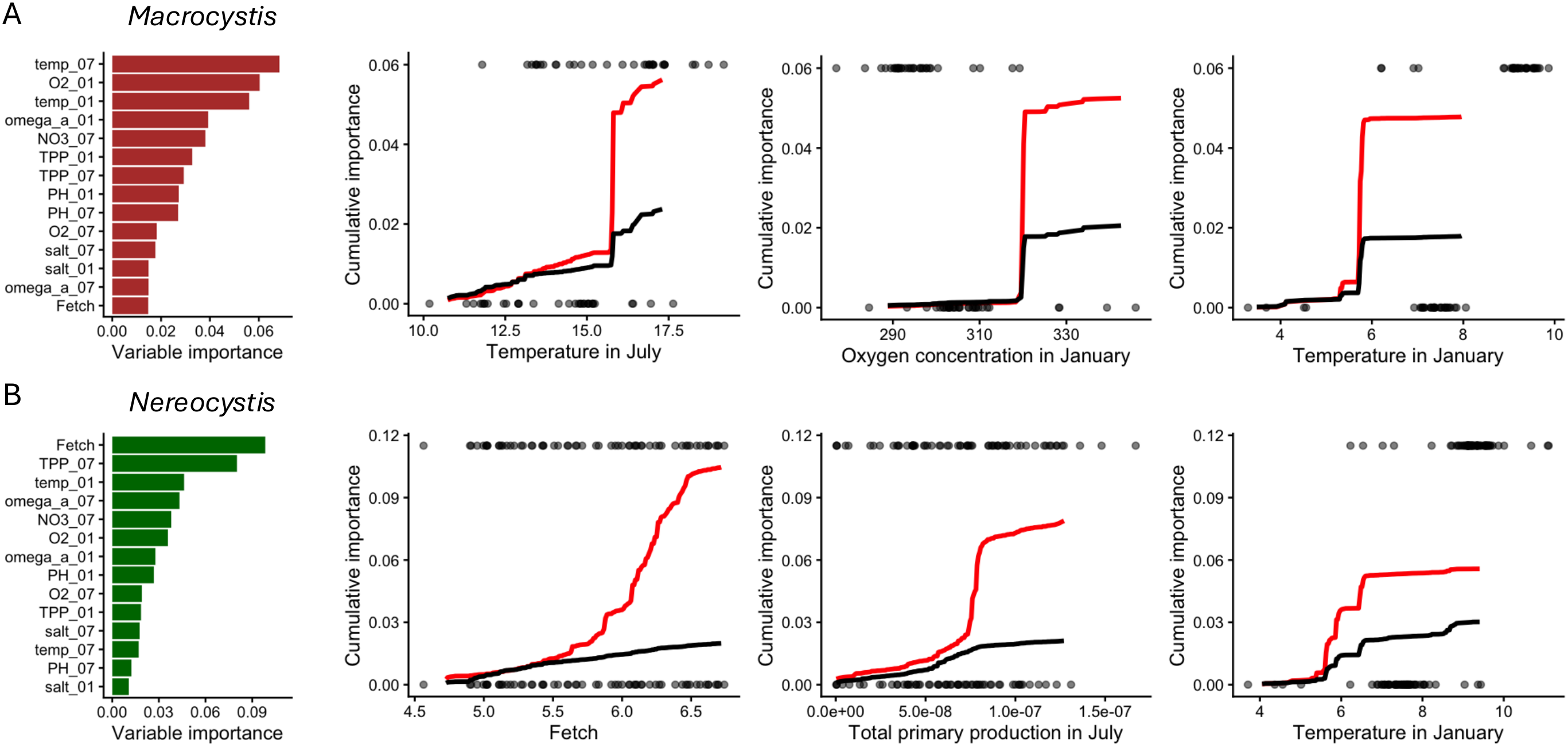
Genetic turnover along environmental gradients. Relative importance of environmental variables in the gradient forest models and genomic turnover for adaptive (red line) and neutral (black line) SNPs across environmental gradients. For each species, the top three environmental variables are shown. Circles correspond to environmental values for sampled sites in historical (bottom) and future (RCP.4.5; top) environments.

### Genomic offsets to predict vulnerability to climate change

First, we used GF analysis to determine the relative importance of environmental variables shaping the distribution of adaptive genetic variation in each of the species. For *Macrocystis*, temp_07, O_2__01, and temp_01 were the most relevant environmental variables (Fig. 3), with all three showing a strong and rapid allele frequency turnover (Fig. 3A; Fig. S2). For *Nereocystis*, fetch was the most relevant environmental variable, followed by total primary production in July (TPP_07), and temp_01 (Fig. 3B; Fig. S2).

Genomic offsets (GOs) were larger for the warming scenario with no mitigation (RCP8.5) than the warming scenario with moderate mitigation (RCP4.5), though both estimates were strongly correlated (r = 0.97 and 0.91 for *Macrocystis* and *Nereocystis*, respectively), so we focus below on results of the moderate mitigation scenario (RCP4.5) to be conservative. In general, warmer sites during summer showed larger GOs than colder ones in both species (correlation between local GO and temp_07 (r) = 0.57 and 0.31 for *Macrocystis* and *Nereocystis*, respectively). Among regions, the northernmost ones (the inner coast of Haida Gwaii and the North Coast; see Fig. 1A for definitions of regions) and Barkley Sound showed the highest values of Local GOs, indicating that these regions may be the most vulnerable to global change (Fig. 4A). In addition, both Population GOs and Location GOs were significantly lower than Local GOs (Fig. 4B), indicating that migration can significantly attenuate mismatch to future environments. This was also true when either 50 km (Population GO_50km_ and Location GO_50km_) or within-region (Population GO_REGION_ and Location GO_REGION_) migration restriction was employed, although attenuation was stronger when migration was allowed within regions (Fig. 4B).

**Figure 4.**
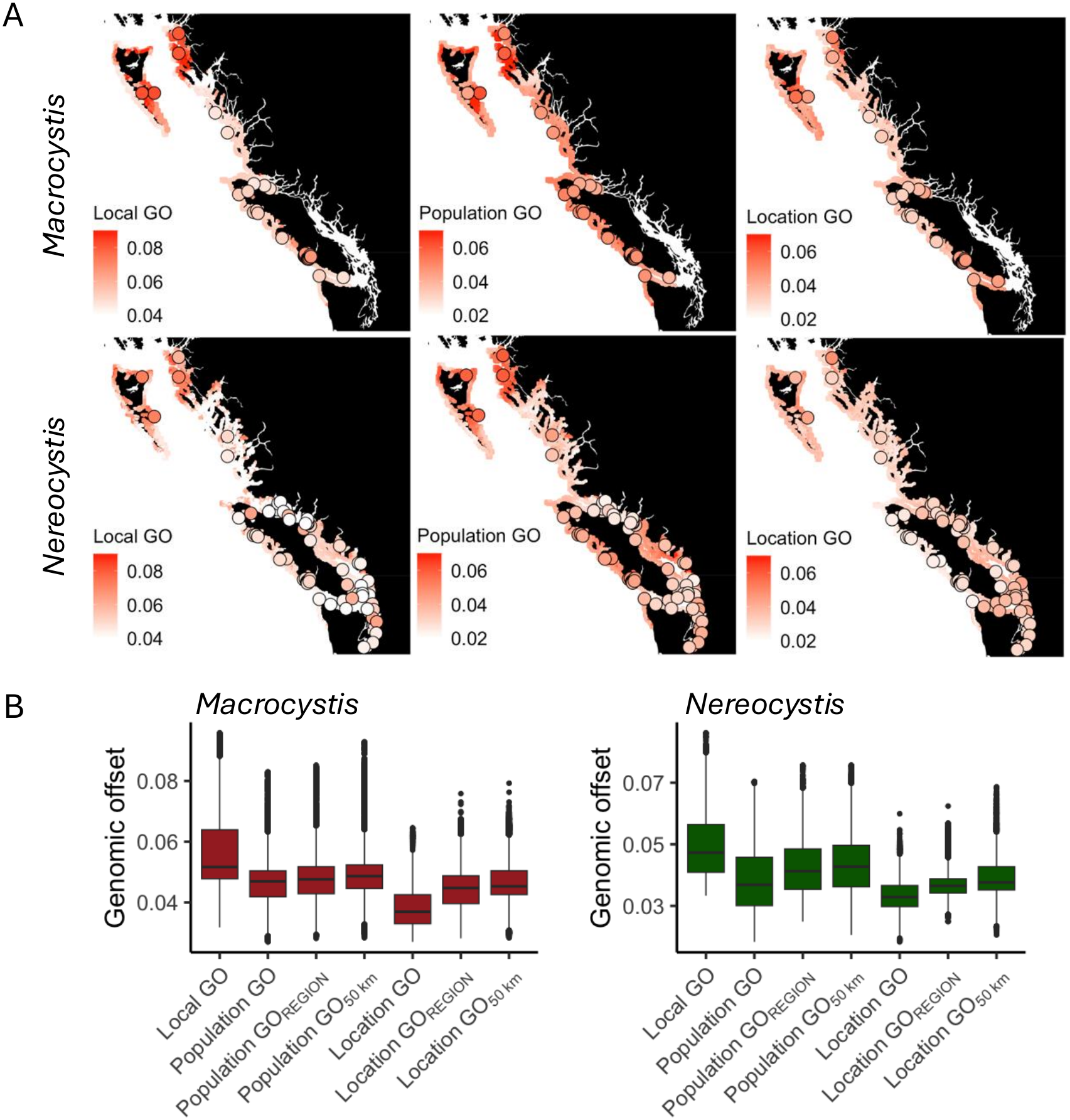
Genomic vulnerability to climate change. A: Geographic distribution of genomic offsets for *Macrocystis* (top panels) and *Nereocystis* (bottom panels). Open circles represent genotyped populations used to build the models. B: Genomic offsets estimated for grid data.

As migration is predicted to attenuate maladaptation to projected environments, for each species, we mapped the geographic distance between each site and its predicted optimal source population under future environments (Fig. 5). For *Macrocystis*, the Strait of Juan De Fuca, Haida Gwaii and the North Coast showed the lowest distances, indicating that local or nearby genotypes are the best option under future environments, regardless of whether estimated GOs are overall low (Strait of Juan De Fuca) or high (Haida Gwaii and North Coast). For all the other regions (i.e., North Vancouver Island, Outer South Coast, Central Coast, and Barkley Sound), long-distance migrations are required to maintain optimal genotype-environment combinations under future environmental conditions. For *Nereocystis*, the Strait of Juan de Fuca and North Vancouver Island showed the lowest distance values, while the remaining regions showed overall long distances between sites and their predicted optimal source population under future environments (Fig. 5).

**Figure 5.**
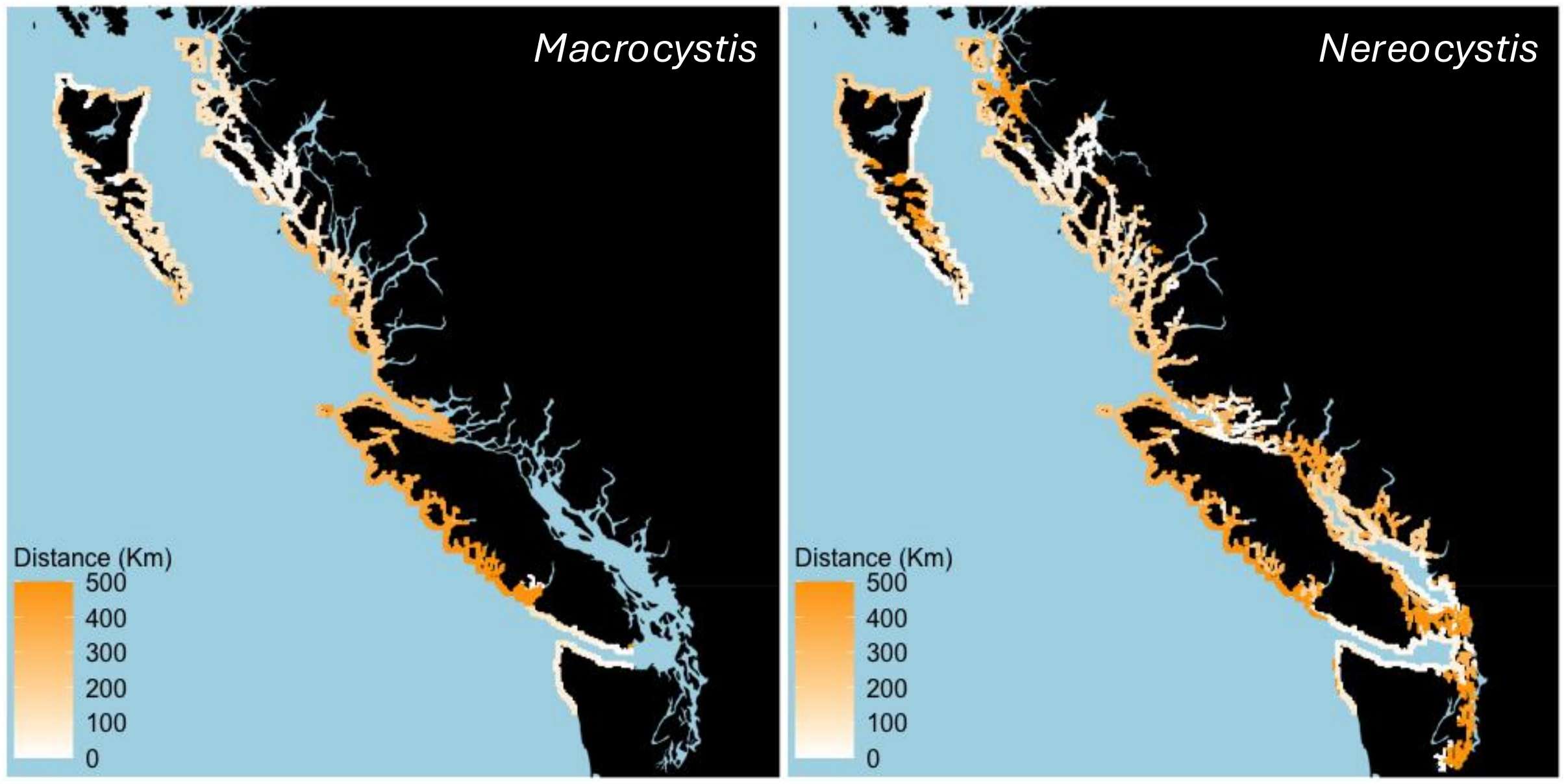
Geographic distance between each recipient population and its optimal source under future climates for *Macrocystis* (left) and *Nereocystis* (right).

### Genomic offsets predict local kelp declines

Our models used presence-absence data for two time periods (historical and ca. 2020) to estimate risk of extirpation as the probability that a site historically containing kelp would no longer (ca. 2020) contain kelp given climate change to date. For both species, we found a significant association, in the predicted direction, between genomic offset for 2020 and the risk of extirpation (Fig. 6A), though model fit was substantially better for *Macrocystis* (p < 2 x 10^-16^; McFadden-pseudo-r^2^ = 0.63) than for *Nereocystis* (p = 0.002; McFadden-pseudo-r^2^ = 0.03). Similar results were found when climate data for 2020 were interpolated using the no mitigation scenario RCP8.5 (McFadden-pseudo-r^2^ = 0.58 and 0.01 for *Macrocystis* and *Nereocystis*, respectively), and when genomic offsets were recalculated for *Nereocystis* excluding fetch (McFadden-pseudo-r^2^ = 0.02). For *Macrocystis*, the model with genomic offset as the predictor variable outperformed the null model with temperature in July as predictor (AIC = 111.7 vs. 146.1; r^2^ = 0.63 vs. 0.49) while the opposite was found for *Nereocystis* (AIC = 350.5 vs. 292; r^2^ = 0.03 vs. 0.21). In addition, none of the environmental variables in *Nereocystis* models correlated strongly with the risk of extirpation (Table S2).

**Figure 6.**
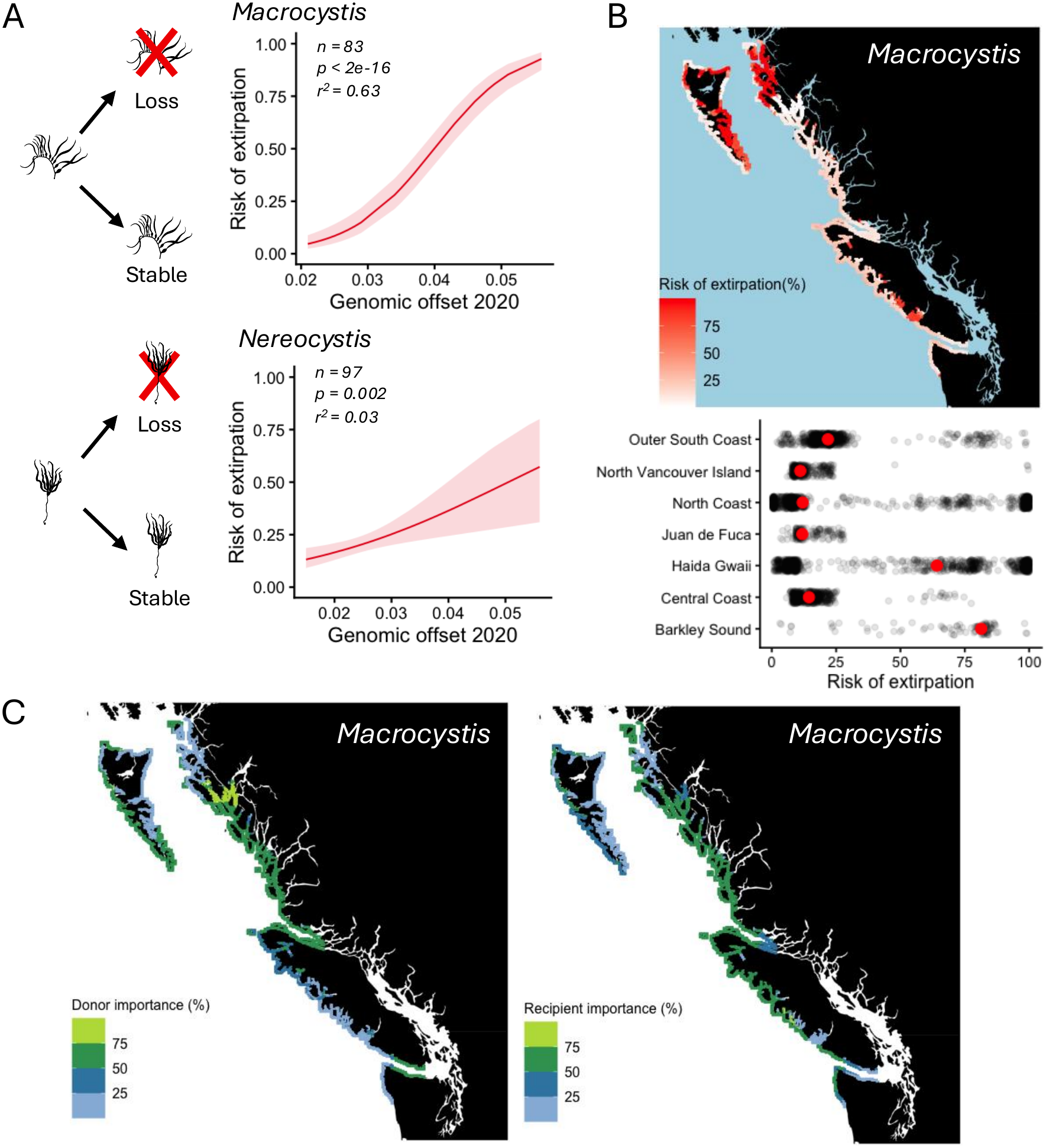
Genomic offsets predict local kelp declines. A: Weighted logistic model between genomic offsets for 2020 and the risk of extirpation. B: Top: map of the risk of extirpation by 2040 for *Macrocystis* under the mitigation scenario (RCP.4.5). Bottom: predicted risk of extirpation by region with red dots indicating median values. C: Donor and recipient importance (%) for *Macrocystis*.

For *Macrocystis*, we found that without migration, some regions (Barkley Sound, Haida Gwaii, and the northernmost North Coast) show an overall high risk of extirpation (Fig. 6B), suggesting high vulnerability to climate change. Threshold-based statistics further suggest challenges for these locations. Areas with the highest risk of extirpation had low (< 25%) DI, suggesting there are few suitable locations in the future for current populations, and low (< 25%) RI, suggesting that future environmental conditions at these locations are suitable only for few current populations. In contrast, for areas with the lowest risk of extirpation (North Vancouver Island and the Central Coast), both DI and RI were relatively high (> 50%) (Fig. 6C).

## DISCUSSION

We characterized genomic patterns of environmental adaptation in two canopy-forming kelp species (*Macrocystis tenuifolia* and *Nereocystis luetkeana*) and demonstrated that many kelp populations are at risk of extirpation in the coming decades, but this risk could be reduced through assisted migration of pre-adapted genotypes to match future climates. Despite largely co-occurring across BC and Washington, the species differed in the environmental gradients that showed the strongest signatures of adaptation. Geographic sites and regions differed substantially in their predicted vulnerability to future climates and in their geographic distance to the source population with the optimal genomic match to future climates (i.e., lowest genomic offset). For *Macrocystis*, but not for *Nereocystis*, genomic offsets strongly predicted kelp loss observed to date, thus validating the link between inferred genomic vulnerability and actual ecological outcomes and allowing us to more confidently use genomic offsets to predict the risk of extirpation in future decades. For *Nereocystis*, no environmental variables correlated strongly with observed declines, suggesting that unmodelled non-environmental factors (e.g., biotic interactions) may be relevant to *Nereocystis* persistence. We also explored several possible assisted migration management scenarios and found that current conservative policies for kelp movement do allow scope for assisted migration to reduce extirpation risk, but that longer distance migration is needed to substantially reduce extirpation risk for many populations. Overall, our results confirm the ecological relevance of genomic offsets and make predictions about the effects of different management strategies and which populations are most vulnerable, demonstrating the utility of genomic offsets to guiding kelp conservation strategies in the face of global change.

### Kelp populations show signatures of local adaptation to environmental gradients

Because predictions of genomic offsets rely on historical genotype-environment associations, accurately identifying genomic regions underlying environmental adaptation is critical for predicting response to climate change. Most studies in kelp have used reduced representation sequencing methods (e.g., Vranken et al. 2021; Wood et al. 2021; Minne et al., 2025) or sampled across small geographic areas (Abbot et al., 2026), thus missing many adaptive regions due to low genetic and/or spatial resolution (Lowry et al., 2017; Layton et al., 2024). Our study addresses those main caveats by combining whole genome sequencing, a comprehensive sampling across the Northeast Pacific Coastline, and high-resolution climate data (both present and future).

We identified the most relevant environmental drivers of local adaptation, and several genomic regions strongly associated with environmental gradients. Interestingly, even when species distributions largely overlap across most of the studied area, the most relevant environmental drivers of local adaptation were distinct. For *Macrocystis*, the mean temperature in July showed the most rapid and strongest genetic turnover, but it had little importance for *Nereocystis*, while the opposite was true for fetch, a proxy for wave exposure. The importance of fetch for *Nereocystis* but not *Macrocystis* adaptation may reflect the wider distribution of *Nereocystis* in BC (Druehl 1978). *Nereocystis* has a higher tolerance than *Macrocystis* to low salinity, allowing it to occupy both exposed outer coasts as well as inner seas and narrow fjords. Consequently, *Nereocystis* occupies a wider gradient in fetch, perhaps making it a more important axis of adaptation overall.

It is more surprising that the mean temperature in July was important for adaptation in *Macrocystis* but not *Nereocystis*, given that extreme summer temperatures have been linked to die-off in both species (Rogers-Bennett and Catton 2019; Mora-Soto et al., 2024; Starko et al., 2024). However, because *Nereocystis* has historically been capable of occupying some of the warmest waters in BC, including the north Salish Sea (Mora-Soto et al. 2024), it is possible that summer temperatures have not historically been highly stressful for *Nereocystis* and were not a primary axis of adaptive population differentiation. Though fetch and summer temperature are known to be relevant for kelp distribution and persistence (Pfister et al., 2018; Mora-Soto et al., 2024; Starko et al., 2024; Man et al., 2025), to our knowledge, this is the first study showing different adaptive responses between co-occurring kelp species in the Northeast Pacific Coastline. This information is critical for designing management zones based on environmental factors and predicting species’ vulnerability to global change (Yu et al., 2022; Lachmuth et al., 2023; Jacquemart et al., 2025).

### Genomic offsets and importance for conservation

The identification of regions most vulnerable to global change is key to prioritizing conservation and restoration efforts (Theissinger et al., 2023; Timm et al., 2023; Wood et al., 2024b). In our study, genomic offsets differed greatly among regions (though also within regions), with warmer sites showing greater offsets than colder ones. Similar patterns were observed in other kelp species and regions, e.g., *Ecklonia radiata* in Australia (Vranken et al., 2021; Minne et al., 2025). However, unlike other regions where ocean temperature (and predicted genomic offsets) follows a latitudinal gradient, in the Northeast Pacific, the complex geography of the coastline creates a mosaic of cold and warm microclimates across the latitudinal gradient, resulting in mosaics of differential predicted vulnerability to global change. Supporting this, large differences in kelp abundance are associated with thermal gradients within distances as short as 16 km (Starko et al., 2022). As historical temperatures at warm edges represent the extreme of species’ tolerance, relatively small temperature increases can make these environments unsuitable for the species (Rehm et al., 2015; Fredston-Hermann et al., 2020; Vranken et al., 2021; Minne et al., 2025). For this same reason, populations from the warmest sites are more susceptible to marine heat waves (Tait et al., 2021; Starko et al., 2022). The effects of marine heat waves are not included in our models and will likely exacerbate populations’ vulnerability to global change (Rogers-Bennett and Catton 2019; Tait et al. 2021). Incorporating the effects of marine heatwaves (i.e., modelled or observed temperature extremes) could also potentially improve model performance, if temperature extremes are a factor strongly influencing patterns of local adaptation.

We identified the northernmost region, which includes the inner coast of Haida Gwaii and the northernmost North Coast, as the most vulnerable to global change. Declines in kelp abundance have been observed in these regions (Starko et al. 2024), especially at the warmest sites (Gendall et al. 2025), suggesting a direct link between kelp abundance and ocean warming (Starko et al., 2024; Gendall et al., 2025). Though this pattern is counter to the current paradigm that low latitude populations are most at risk, ocean temperature – the best predictor of genomic offsets in our study – did not follow a latitudinal gradient along the Pacific Coastline, which is characterized by high microclimate variation.

Besides the northernmost regions, other regions like Barkley Sound for *Macrocystis* and the Strait of Georgia for *Nereocystis* also showed high genomic offsets, which is in line with previous local-scale studies showing declines in kelp abundance in these regions (Mora-Soto et al., 2024; Starko et al., 2024). In contrast, regions like North Vancouver Island, the Strait of Juan de Fuca, and the Central Coast of BC showed the lowest genomic offsets, suggesting kelps are overall resilient in these areas, which is supported by local-scale studies showing that kelp abundance has been stable in these regions (Mora-Soto et al., 2024; Man et al., 2025). Therefore, our study identified vulnerable regions where management interventions should be prioritized and helps explain contrasting patterns of kelp vulnerability/resilience observed in local-scale studies. The qualitative agreement between regions predicted to be most vulnerable and those that have exhibited recent declines suggests that kelp populations in the most vulnerable areas may already be facing maladaptation, highlighting the need for flexible kelp management strategies that consider local adaptation.

Validation of genomic offset predictions are scarce, and they are biased towards plant species that can be grown in common gardens (Fitzpatrick et al., 2021; Luo et al., 2025; Verrico et al., 2026; reviewed in Fitzpatrick et al., 2026). In marine systems, validations of genomic offsets are even scarcer (Layton et al., 2024; but see Selmoni et al., 2025), which has hindered the implementation of this tool in marine conservation programs. Here, using kelp distribution data obtained mostly from aerial images, we found the expected positive association between genomic offsets and the risk of extirpation in both species, thus validating genomic predictions. However, validation models performed substantially better for *Macrocystis* than for *Nereocystis*. Lind & Lotterhos (2025) evaluated the performance of genomic offsets across a simulated data set and found that genomic offsets perform best when local adaptation is strong, drivers of adaptation are known, and future environments are not extremely novel relative to current ones. In our study, local adaptation seems to be weaker in *Nereocystis* than in *Macrocystis* (lower isolation by environment; Fig. 2C), and no environmental variables correlated strongly with the risk of extirpation in *Nereocystis* (Table S2), suggesting that unmodelled environmental factors contributed to *Nereocystis* persistence.

Among these non-environmental factors, overgrazing by herbivores (mainly sea urchins) is likely the most relevant, affecting kelp persistence across its range (Filbee-Dexter and Scheibling, 2014; Ling et al., 2015; Starko et al., 2022). Though sea urchins consume both *Macrocystis* and *Nereocystis,* as they prefer young kelp recruits, the negative impact of overgrazing could be strongest for the annual species *Nereocystis*. We also speculate that local extirpation and recolonization dynamics (e.g., due to stochastic population processes or inter-annual environmental variation) may be more variable for the annual species *Nereocystis* than longer-lived *Macrocystis* and could potentially explain the poorer model fit for *Nereocystis.* Another factor likely affecting validation models is the time lapse between observations. The median time lapse is 14 years, for most observations starting in 2004 – 2007, thus missing sites where extirpation occurred before the beginning of the study period. Also, the time lapse was originally thought to capture the effects of a sustained heat event during 2014 – 2016 on kelp persistence (Starko et al., 2024), and according to our models, summer temperatures are more relevant for *Macrocystis* than for *Nereocystis*. We also acknowledge that sharp transitions in temperature between the northernmost populations (from Haida Gwaii and North Coast) and the rest may be the result of a strong covariance between population structure and environmental variation and affect genomic offset estimations, as populations crossing steep transitions will have the largest genomic offsets.

Including more sites covering a broader area, longer timeseries, the frequency and intensity of extreme weather events (e.g., heatwaves), and proxies for both biotic interactions and human pressure is expected to improve our models and to better inform kelp forest conservation and restoration. To this end, remote sensing technologies could represent a powerful, high-throughput, and cost and time-efficient method to further validate genomic predictions in kelp. However, if biotic factors such as herbivore grazing pressure play a more important role than abiotic environmental variables in driving local population loss – as we have speculated may be the case for *Nereocystis* – then the practical utility of genomic offset tools in guiding local kelp conservation and restoration may be limited.

### Complementary offset statistics identify valuable germplasm for restoration

Restoration efforts in kelp are becoming important in several regions of the globe (Eger et al., 2022; Wood et al., 2024b) and promising approaches for large-scale restoration are under development (Fredriksen et al., 2020; Dykman et al., 2025). Yet, there are no clear guidelines for the optimal source population selection, which is an important factor determining restoration success (Houde et al., 2015). Here, we show that local genotypes are rarely the best adapted under projected future environmental conditions and long-distance migrations among regions are often required to minimize genotype-environment mismatch. In BC and Washington, the 50 km rule restricts the transfer of kelp between populations more than 50 km apart (Cui 2023; McConnell et al. 2024), narrowing down the options for the optimal source population selection. Our results show that moving genotypes either within 50 km or within geographic regions (largely based on neutral genetic structure) both offer significant and similar attenuation of extirpation risk, even though geographic regions as defined here often extend much further than 50 km. This finding likely reflects the high microclimatic variation along the coasts of BC and Washington, where geographically proximate locations can experience dramatically different environmental conditions (but see Fig. 1D).

One caveat to the single best-match approach (i.e., Population GO and Location GO) is that it uses all possible kelp habitats as a possible donor, but neither kelp species occupies all possible locations in its range. This means that we may predict an ideal donor population to exist at a location where there is no current kelp. However, at least for *Macrocystis*, by using threshold-informed metrics (i.e., DI and RI; Fig. 6B), we showed that populations and locations with low Population/Location GO also showed high DI and RI (i.e., multiple suitable combinations of populations and locations) and vice versa (Fig. 6C). Threshold-informed metrics also provide more flexibility for management in conservation, as potential source populations can be selected from a pool of suitable donors rather than a single best-match source. Although single best-match and threshold-informed metrics can be complemented to inform management decisions in *Macrocystis*, we acknowledge that for *Nereocystis*, more robust models linking genomic offsets with extirpation are needed before genomic offset thresholds could be reliably used to select source populations for restoration.

Another caveat to our approach relates to the fundamental assumption of local adaptation to microclimates in genomic offset analyses. These analyses assume the existence of adaptive loci whose allele frequencies are correlated with environmental variation, yet relatively little is known about the extent to which kelp populations are locally adapted in BC and Washington (Bemmels and Owens, 2026). Local adaptation is laborious to study with field-based reciprocal transplants in kelp and existing field studies from this region over small spatial scales have often failed to find evidence of local adaptation (Fales et al., 2023; Weigel et al., 2023; Dykman et al., 2025) or have found mixed results (Dykman et al., 2026), though local adaptation has been demonstrated in *Macrocystis* from other regions of the world (reviewed in Bemmels and Owens, 2026) and inferred from genomic data in *Nereocystis* from Puget Sound (Abbott et al., 2026). Weak or nonexistent local adaptation could plausibly be a contributor to poor ability to explain observed declines in *Nereocystis.* However, even if kelp populations exhibit local adaptation overall across broad environmental gradients, this does not necessarily mean that every population is optimally locally adapted. For example, a kelp population that is present in a given microclimate without necessarily being locally adapted to that microclimate could be selected as an optimal donor in our models but would in reality provide little benefit (Dykman et al., 2025). A potential solution to overcome this caveat is the characterization of the adaptive genetic variation of potential donor populations, i.e., testing if donors indeed represent novel genotypes predicted to be well adapted to an outplanting environment. Though whole-genome sequencing for many samples may be cost-prohibited, our study provides a low number of genomic regions (< 100 per species) strongly associated with environmental gradients, which could be target-sequenced to allow cost-effective genotyping. Combining target sequencing with pool-seq approaches would further reduce genotyping costs, facilitating its adoption in conservation programs.

Genomic offsets and extirpation risk are lowest of all when there are no restrictions placed on migration distance. This suggests that extreme long-distance migration has the potential to substantially improve resilience in most regions. The potential adaptive benefits of long-distance assisted migration should be weighed against the possibility of outbreeding depression (Frankham et al. 2011) between groups that may have been genetically isolated for long periods of time (Bemmels et al., 2026). In addition, our models do not consider the possibility that populations may be able to adaptively evolve (i.e., experience adaptive change in allele frequencies), reducing the need for long-distance migration, although recent declines in the Northeast Pacific Coastline have been linked to ocean warming (Starko et al., 2022; Gendall et al., 2025) suggesting that kelp species are not adapting fast enough to keep pace with global change. Also, long-distance migrations may negatively alter biotic interactions of kelp (e.g., with urchins or coralline algae), which are not included in our models but are potentially relevant for local adaptation (Twist et al., 2024). Finally, we also acknowledge that deliberately altering the genetic composition of natural populations raises bioethical issues that would require careful consideration from stakeholders before implementation (Coleman et al. 2020). Despite potential side-effects and uncertainties, long-distance migration may be the only viable option for restoring areas where kelp have long been extirpated, and nearby populations either do not exist or are declining.

Regardless of the management policy pursued for the transport of individuals, we identified geographic areas that are the most likely to exhibit genomic mismatch with future climates. These sites are mostly distributed in Haida Gwaii and the North Coast for both species, and additionally in Barkley Sound for *Macrocystis*. Many sites within these regions are still predicted to have a genomic mismatch (and predicted risk of extirpation in *Macrocystis*) under most migration management scenarios, suggesting that even targeted interventions may be insufficient to protect some kelp populations in BC from extirpation. Importantly, northern regions harbour the highest genetic diversity within BC and Washington for both species (Bemmels et al. 2025), suggesting that they may be important reservoirs for genetic diversity in general, potentially including adaptive variation to factors other than climate. This combination of factors (high risk and high genetic diversity) highlights that Haida Gwaii and the North Coast should be a priority for conservation resources and research about the impacts of global change. In contrast, areas predicted to have a lower mismatch with future climates (such as North Vancouver Island and Juan de Fuca) could be ideal candidate locations for marine protected areas or other tools emphasizing *in situ* conservation.

## CONCLUSIONS

Genomic offsets are becoming an increasingly important tool to predict maladaptation driven by global change. However, the lack of validation of such predictions has prevented the widespread implementation of GEA analyses in marine and terrestrial conservation programs. Here, using a seascape genomics approach in two kelp species, we identified areas along the Northeast Pacific Coastline of high genomic mismatch to future climate that should be prioritized in conservation planning. For one of the species, *Macrocystis*, we also link genomic offsets to extirpation risk, which allows us to quantify the predicted benefit of assisted migration in ecologically meaningful terms (i.e., risk of extirpation). The link between genomic offsets and extirpation observed to date further suggests that the most vulnerable populations may already be facing maladaptation to changing climates, highlighting the need for intervention to ensure their persistence in changing environments. Our results can inform the selection of source populations for assisted migration or restoration in regions where local populations have been extirpated and can help policymakers calculate the costs and benefits of assisted migration policy. Nonetheless, further validation (e.g., of transplanted populations) and incorporation of non-environmental factors potentially affecting kelp persistence are needed to improve model predictions and to better inform conservation and restoration, especially for *Nereocystis*.

## ACKNOWLEDGEMENTS

We thank the Gitga’at, Gitxaała, Haida, Haisla, Heiltsuk, Kitasoo-Xai’xais, Kitselas, Kitsumkalum, K’ómoks, Mamalilikulla, Metlakatla, Tlowitsis, and Wei Wai Kum First Nations for permission to reanalyze DNA sequences from samples collected from their territories. For a Biocultural Notice of cultural rights and responsibilities regarding these samples see Bemmels et al. (2025). Funding was provided by the Natural Sciences and Engineering Research Council of Canada (RGPIN-2021-02482 to GLO) and Genome British Columbia (GIRAFF Grant to GLO and LHR). Postdoctoral support was provided by a Mitacs Accelerate Postdoctoral Fellowship to JBB and a Biodiversity Research Centre Fellowship to FH.

## Supplementary materials

**Figure S1.**
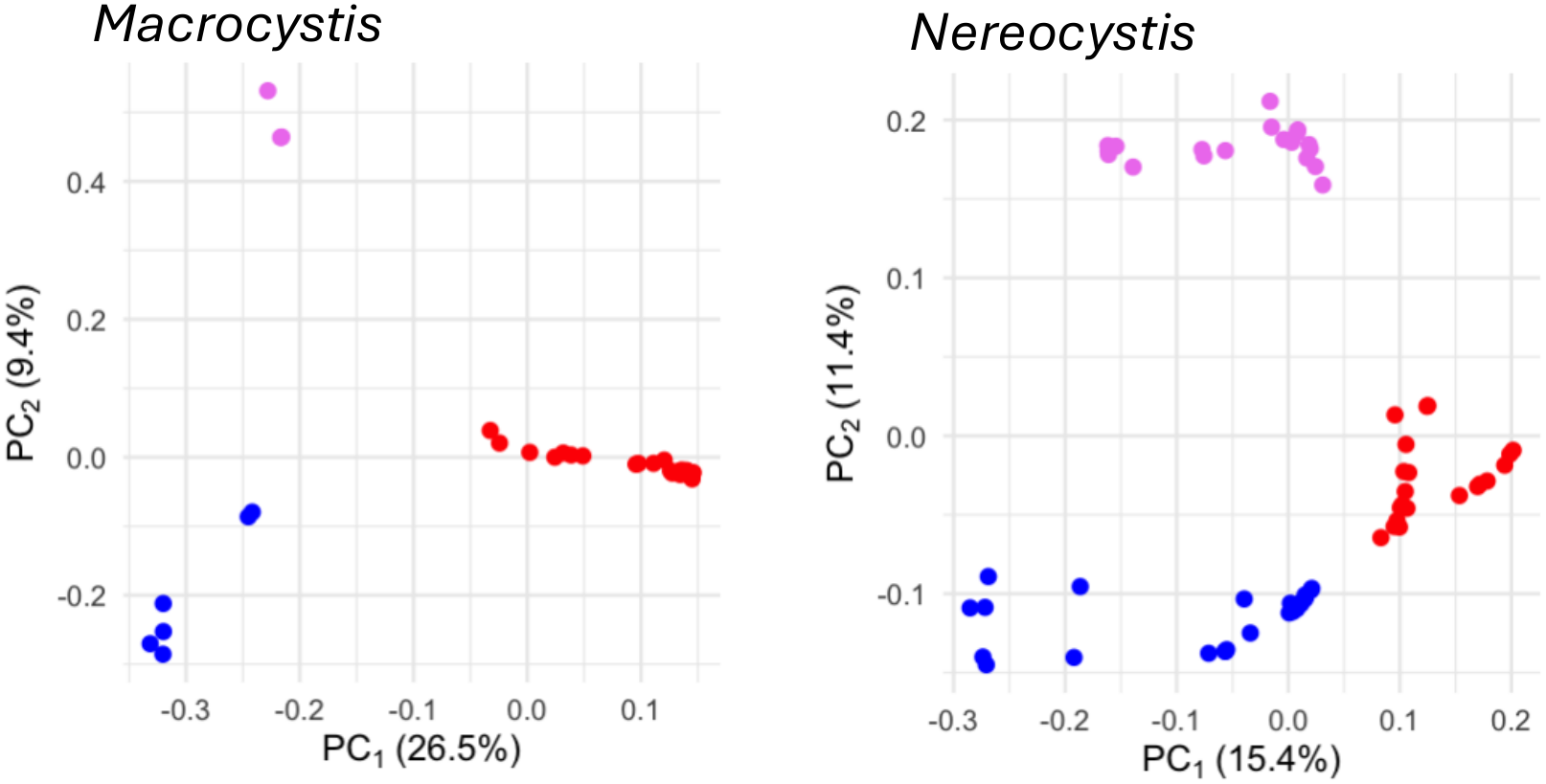
Genetic structure for *Macrocystis* (left) and *Nereocystis* (right) using 43,933 and 116,707 SNPs in low linkage disequilibrium. Each dot represents a population (site) with at least three individuals. Putative ancestral genetic clusters are shown in different colours.

**Figure S2.**
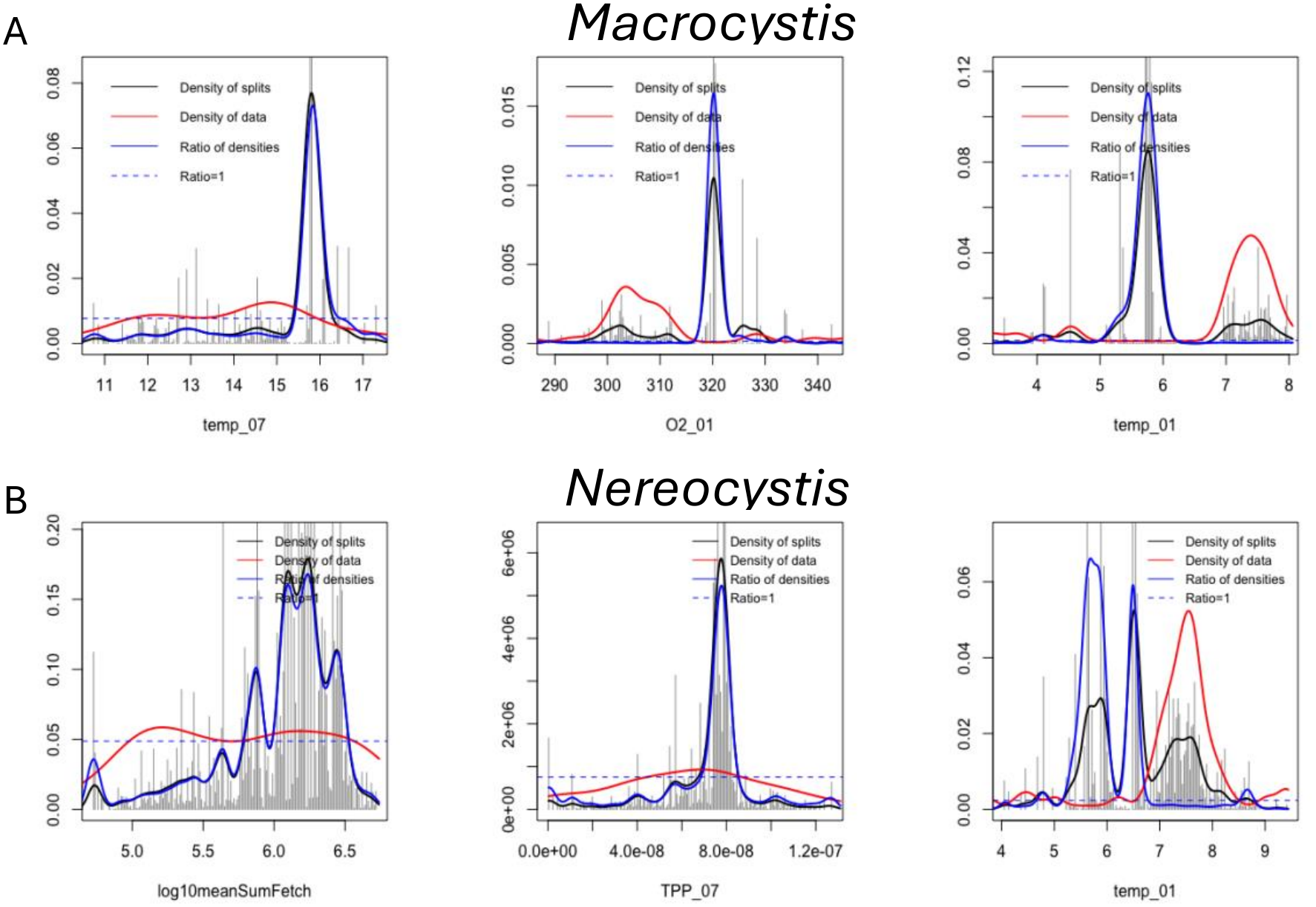
Split-density plots for the three most relevant environmental variables for *Macrocystis* (A) and *Nereocystis* (B).

**Table S1.** Environmental variables used in this study. The final set of variables retained after pairwise correlation < |0.8| is highlighted in bold.

| Variable | Description | Units |
|---|---|---|
| **Fetch** | **log10 mean Sum Fetch** |  |
| Alkalini_01 | Total alkalinity concentration in January | mmol m-3 |
| Alkalini_07 | Total alkalinity concentration in July | mmol m-3 |
| Cflx_01 | Air-sea CO2 flux in January | mol m-2 s-1 |
| Cflx_07 | Air-sea CO2 flux in July | mol m-2 s-1 |
| DIC_01 | Dissolved inorganic concentration in January | mmol m-3 |
| DIC_07 | Dissolved inorganic concentration in July | mmol m-3 |
| maxO2_01 | Maximum oxygen concentration in January | mmol m-3 |
| maxO2_07 | Maximum oxygen concentration in July | mmol m-3 |
| maxPH_01 | Maximum pH in January |  |
| maxPH_07 | Maximum pH in July |  |
| maxtemp_01 | Maximum temperature in January | Deg C |
| maxtemp_07 | Maximum temperature in July | Deg C |
| minO2_01 | Minimum oxygen concentration in January | mmol m-3 |
| minO2_07 | Minimum oxygen concentration in July | mmol m-3 |
| minPH_01 | Minimum pH in January |  |
| minPH_07 | Minimum pH in July |  |
| mintemp_01 | Minimum temperature in January | Deg C |
| mintemp_07 | Minimum temperature in July | Deg C |
| NO3_01 | Nitrate concentration in January | mmol m-3 |
| **NO3_07** | **Nitrate concentration in July** | **mmol m-3** |
| **O2_01** | **Dissolved oxygen concentration in January** | **mmol m-3** |
| **O2_07** | **Dissolved oxygen concentration in July** | **mmol m-3** |
| **omega_a_01** | **Aragonite saturation state in January** |  |
| **omega_a_07** | **Aragonite saturation state in July** |  |
| **PH_01** | **pH in January** |  |
| **PH_07** | **pH in July** |  |
| **salt_01** | **Sea water practical salinity in January** | **PSU** |
| **salt_07** | **Sea water practical salinity in July** | **PSU** |
| sigma_01 | Sigma potential density in January | kg m-3 |
| sigma_07 | Sigma potential density in July | kg m-3 |
| TCHL_01 | Total chlorophyll (diatoms + nano chlorophyl concentration) in January | mg m-3 |
| TCHL_07 | Total chlorophyll (diatoms + nano chlorophyl concentration) in July | mg m-3 |
| **temp_01** | **Mean sea surface temperature in January** | **Deg C** |
| **temp_07** | **Mean sea surface temperature in July** | **Deg C** |
| **TPP_01** | **Total primary production (diatoms + nano) in January** | **mol m-3 s-1** |
| **TPP_07** | **Total primary production (diatoms + nano) in July** | **mol m-3 s-1** |

**Table S2.** Output of weighted logistic models for each environmental variable used as predictor and the risk of extirpation in both species. McFadden-pseudo-r2 was calculated as [1-(LLMODEL/LLNULL)]

| Variable | McFadden-pseudo-r<sup>2</sup> |  |
|---|---|---|
|  | *Nereocystis* | *Macrocystis* |
| Fetch | 0.19 | 0.03 |
| NO<sub>3</sub>_07 | 0.25 | 0.3 |
| O<sub>2</sub>_01 | 0.07 | 0.01 |
| O<sub>2</sub>_07 | 0.03 | 0.1 |
| omega_a_01 | 0.16 | 0.05 |
| omega_a_07 | 0 | 0.1 |
| PH_01 | 0.02 | 0.15 |
| PH_07 | 0.04 | 0.58 |
| salt_01 | 0 | 0.16 |
| salt_07 | 0.1 | 0.65 |
| temp_01 | 0.1 | 0.08 |
| temp_07 | 0.21 | 0.49 |
| TPP_01 | 0.03 | 0.39 |
| TPP_07 | 0.25 | 0.02 |
| Genomic offset | 0.03 | 0.63 |

